# Computation of condition-dependent proteome allocation reveals variability in the macro and micro nutrient requirements for growth

**DOI:** 10.1101/2020.03.23.003236

**Authors:** Colton J. Lloyd, Jonathan Monk, Laurence Yang, Ali Ebrahim, Bernhard O. Palsson

## Abstract

Sustaining a robust metabolic network requires a balanced and fully functioning proteome. In addition to amino acids, many enzymes require cofactors (coenzymes and engrafted prosthetic groups) to function properly. Extensively validated genome-scale models of metabolism and gene expression (ME-models) have the unique ability to compute an optimal proteome composition underlying a metabolic phenotype, including the provision of all required cofactors. Here we use the ME-model for *Escherichia coli* K-12 MG1655 to computationally examine how environmental conditions change the proteome and its accompanying cofactor usage. We found that: (1) The cofactor requirements computed by the ME model mostly agree with the standard biomass objective function used in models of metabolism alone (M models); (2) ME-model computations reveal non-intuitive variability in cofactor use under different growth conditions; (3) An analysis of ME-model predicted protein use in aerobic and anaerobic conditions suggests an enrichment in the use of prebiotic amino acids in the proteins used to sustain anaerobic growth (4) The ME-model could describe how limitation in key protein components affect the metabolic state of *E. coli.* Genome-scale models have thus reached a level of sophistication where they reveal intricate properties of functional proteomes and how they support different *E. coli* lifestyles.

## Introduction

An established approach for studying the metabolic capabilities of an organism is the use of genome-scale metabolic models (M-model). M-models have shown significant success in predicting the metabolic capabilities of a cell by integrating all of the experimentally determined enzymatic reactions taking place in an organism [1–4]. These predictions are based on the stoichiometric constraints of the organism’s metabolic network and its metabolic interactions with the environment. Additionally, M-models rely on an empirically derived biomass objective function which relates the growth rate of the simulation to the biosynthesis of all major biosynthetic building blocks needed to synthesize RNA, protein, and other macromolecules [5].

The use of this biomass objective function, however, implies that the abundance of all major components in a cell does not change based on growth rate or condition. In actuality, the macromolecular composition of a cell is highly dependent on its specific growth environment. This variability is due to the fact that the macromolecular composition of a cell is a function of the specific collection of proteins used to sustain growth in a particular environment. A key component of synthesizing a functional proteome--along with translating the proper amino acid sequences and folding the proteins into the proper 3D structure--involves equipping enzymes with the necessary prosthetic groups and coenzymes. These accessory enzyme cofactors often drive the chemical conversions at the heart of an enzyme’s activity, making their presence essential for detectable catalytic activity [6,7]. The functions of some cofactors, such as flavins and iron-sulfur clusters, are so essential for core metabolism that their activity can be traced back to the beginning of life [8]. Thus, ensuring that all coenzymes and prosthetic groups are available to enzymes is essential for any robustly growing organism. The scarcity of one or more of the essential micronutrients can have a profound impact on the metabolic state of an organism, such as the disruption in energy metabolism and lactate secretion that is seen in *E. coli* growing in iron-limited stress conditions [9].

Despite their importance in sustaining metabolism, cofactor biosynthesis is not modeled mechanistically in M-models. This is due to the fact that cofactors are either enzyme prosthetic groups and thus have no modeled metabolic function (pyridoxine, biotin, etc.) or can be recycled (NAD, folates, etc.), meaning there is no metabolic process driving their biosynthesis. Thus, cofactors have often been incorporated into the biomass objective function to force the essential biosynthetic activity of enzymes forming these cofactors [5]. The inclusion of cofactors into biomass objective functions has been studied across various bacterial and archaea species providing insight into the essentiality of individual cofactors in prokaryotes [10]. However, even when included in the biomass objective function, a negligible amount of each cofactor is required to be synthesized for growth, causing cofactor synthesis to have little impact on metabolism overall. Furthermore, the specific requirement of the cofactors in the M-model is condition independent. Modeling efforts have been made to assess how the biomass function composition (lipid and amino acid composition) affects metabolic fluxes [11], but a mechanistic model has not been employed to relate cofactor demand to condition-dependent metabolism.

To that end, M-models have been extended to include the synthesis and use of the gene expression machinery to compute the entire metabolic and gene expression proteome [12–14]. These models integrate <u>M</u>etabolism and <u>E</u>xpression on the genome-scale, termed ME-models, and they are capable of explicitly computing over 80% of the proteome by mass in enterobacteria. ME-models enable a wide range of new biological questions to be investigated including direct computations of proteome allocation [15], metabolic pathway usage, and the effects of membrane and volume constraints [13]. Furthermore, their ability to compute the optimal proteome abundances for a given condition makes them ideal for mechanistically integrating transcriptomics and proteomics data. Here we employ the *E. coli* ME-model [16] to examine the relationship between growth condition and cellular biomass composition. This work presents the first effort to comprehensively study the role that essential cofactors play in defining the metabolic capabilities of *E. coli.*

## Results

### Model Development

*E. coli* K-12 MG1655, along with many other microbes, are capable of *de novo* synthesizing the essential cofactors needed for growth, listed in **Table 1**. Thus, the pathways that require these cofactors are included in both the *E. coli* K-12 MG1655 M-model (iJO1366 [17]) and ME-model (iJL1678b [16]). Unique to the ME-model, however, the activity of the prosthetic groups in **Table 1** is also explicitly modeled, as *i*JL1678b includes a mechanistic accounting of all of the components required to produce a functioning proteome (**Figure 1**). The integration of an enzyme and its prosthetic groups means that for a particular enzymatic reaction to carry flux in the model, not only must the amino acids be synthesized in the proper proportions, but enzyme prosthetic groups must also be available. Therefore, the condition-specific synthesis demand of both prosthetic groups and amino acids can be assessed through ME-model computation.

**Figure 1:**
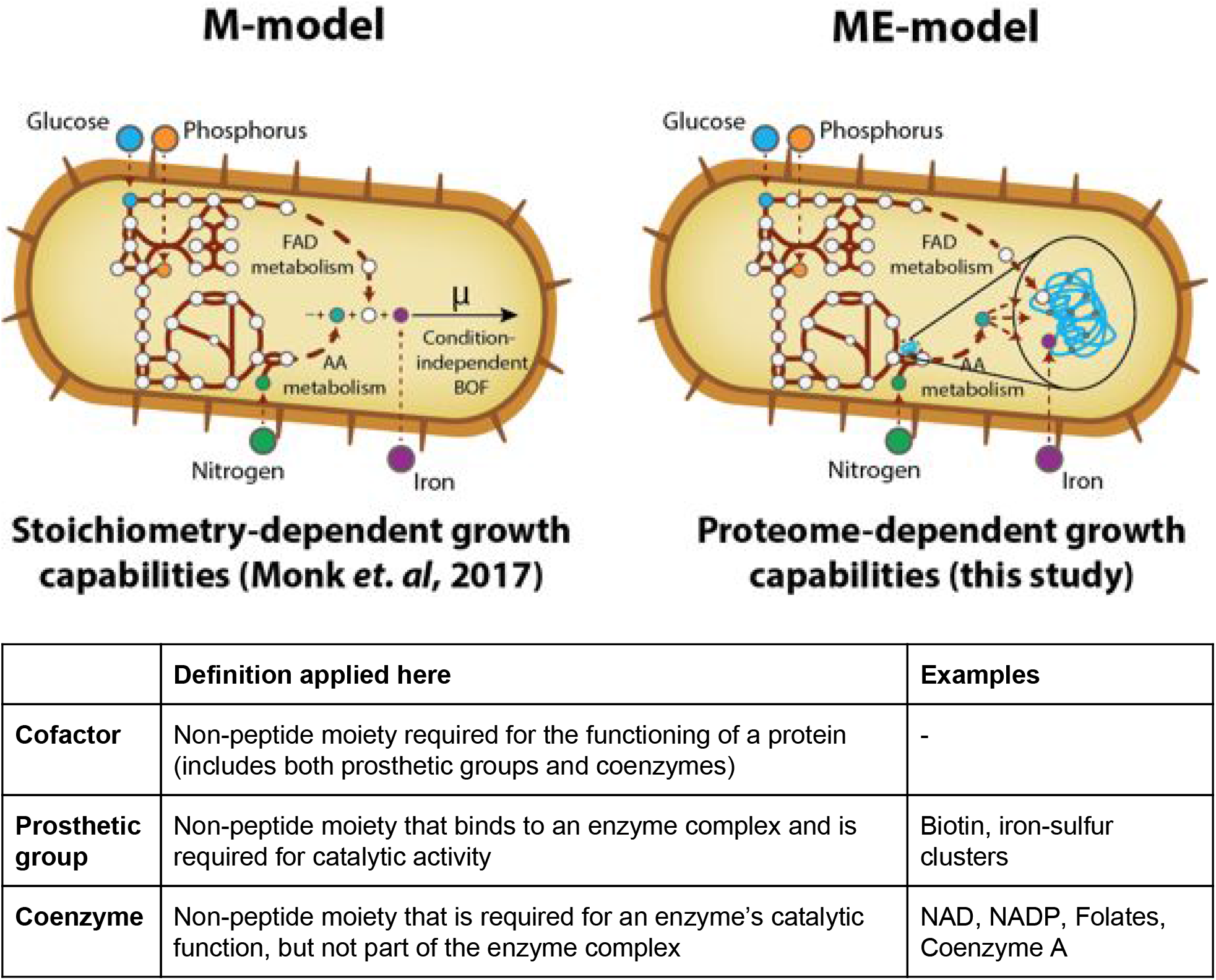
Difference in M- and ME-model scope. M-models offer a means to comprehensively probe the range of enzymatic conversions possible within an organism. This capability is based on the stoichiometry of reactions in the organism’s metabolic network and can be used to predict possible growth supporting nutrient environments [18,19]. By including a mechanistic accounting of enzyme synthesis and activity, ME-models add additional information about the proteome sustaining the growth state. Predictions of the proteome offer the ability to study how proteome allocation and cofactor use affects condition-dependent growth. Due to the inconsistency in how the terms **cofactor, prosthetic group**, and **coenzyme** are used in the scientific literature, the definitions applied in this study are listed in the table.

**Table 1:** Summary of the vitamins synthesized by *E. coli* K-12 MG1655

| Cofactor Name (BIGG ID) | General Function | Cellular Role | Essentiality from Xavier *et al.* | Vitamin |
| --- | --- | --- | --- | --- |
| Thiamin (thm) | Energy metabolism | Prosthetic group | Universal | B1 |
| Riboflavin (ribflv) | Redox metabolism | Prosthetic group/redox coenzyme | Universal (FMN, FAD) | B2 |
| Niacin (nac) | NAD(P) precursor, electron carrier | Coenzyme | Universal (NAD, NADP) | B3 |
| Pantothenic Acid (pnto__R) | CoA precursor, fatty acid biosynthesis | Coenzyme | Universal (CoA) | B5 |
| Pyridoxine (pydxn) | Versatile coenzyme that participates in transamination, decarboxylation, etc. | Prosthetic group | Universal | B6 |
| Biotin (btn) | Required for carboxylase activity | Prosthetic group | Conditional | B7 |
| Folate (thf) | Carrier of single carbon moieties | Coenzyme | Universal | B9 |
| Cobalamin (cbl1) | Certain isomerases and methyltransferases, not essential in *E. coli* K-12 MG1655 | Prosthetic group | Conditional | B12 |
| Menaquinone 8 (mqn8) | Electron carrier | Coenzyme | Conditional (quinones) | K2 |
| Ubiquinone 8 (q8) | Electron carrier | Coenzyme | Conditional (quinones) | - |
| 2-Demethylmenaquinone 8 (2dmmq8) | Electron carrier | Coenzyme | Conditional (quinones) | - |

Unlike prosthetic groups, coenzymes such as NAD and folates act as carriers that donate and accept energy (e.g., electrons carried by NAD(H)) or chemical moieties (e.g., single carbon groups carried by folates). These coenzymes are therefore regenerated throughout the network, meaning the synthesis of coenzymes is not directly coupled to their activity. As is the case in M-models, coenzyme synthesis is not required for growth in the default ME-model and therefore is included as part of a biomass constituent demand reaction [16], analogous to the biomass objective function in M-models. *i*JL1678b was thus modified to account for the activity of these coenzymes and to couple coenzyme synthesis to its metabolic function (see **Methods**). The activity of the coenzymes was determined based on the flux through the reaction that synthesizes the coenzyme (**Table A in S1 Appendix**, see **Methods**). Using this modified ME-model, growth simulations could effectively *de novo* predict the composition of *E. coli* K-12 MG1655’s appropriate biomass objective function in a condition-dependent manner.

### Computing metabolic states and proteome composition including its cofactor needs

These extensions to the ME-model allow us to computationally address a number of important aspects of microbial growth. We present three insightful case studies.

#### (1) Benchmarking ME-model predictions of biomass composition

Following the model extensions outlined above, aerobically and anaerobically computed synthesis fluxes of amino acids, prosthetic groups, and coenzymes were growth normalized to enable a comparison with the *i*JO1366 biomass objective function (**Figure 2**). The computed amino acid synthesis fluxes quantitatively agreed with the empirically derived numbers contained in the *i*JO1366 biomass objective function (BOF) and showed slight differences between aerobic and anaerobic simulations. This agreement suggests that amino acid composition of the proteome under multiple growth conditions is generally well represented by the BOF, though subtle changes in the amino acid composition could be observed depending on the growth condition.

**Figure 2.**
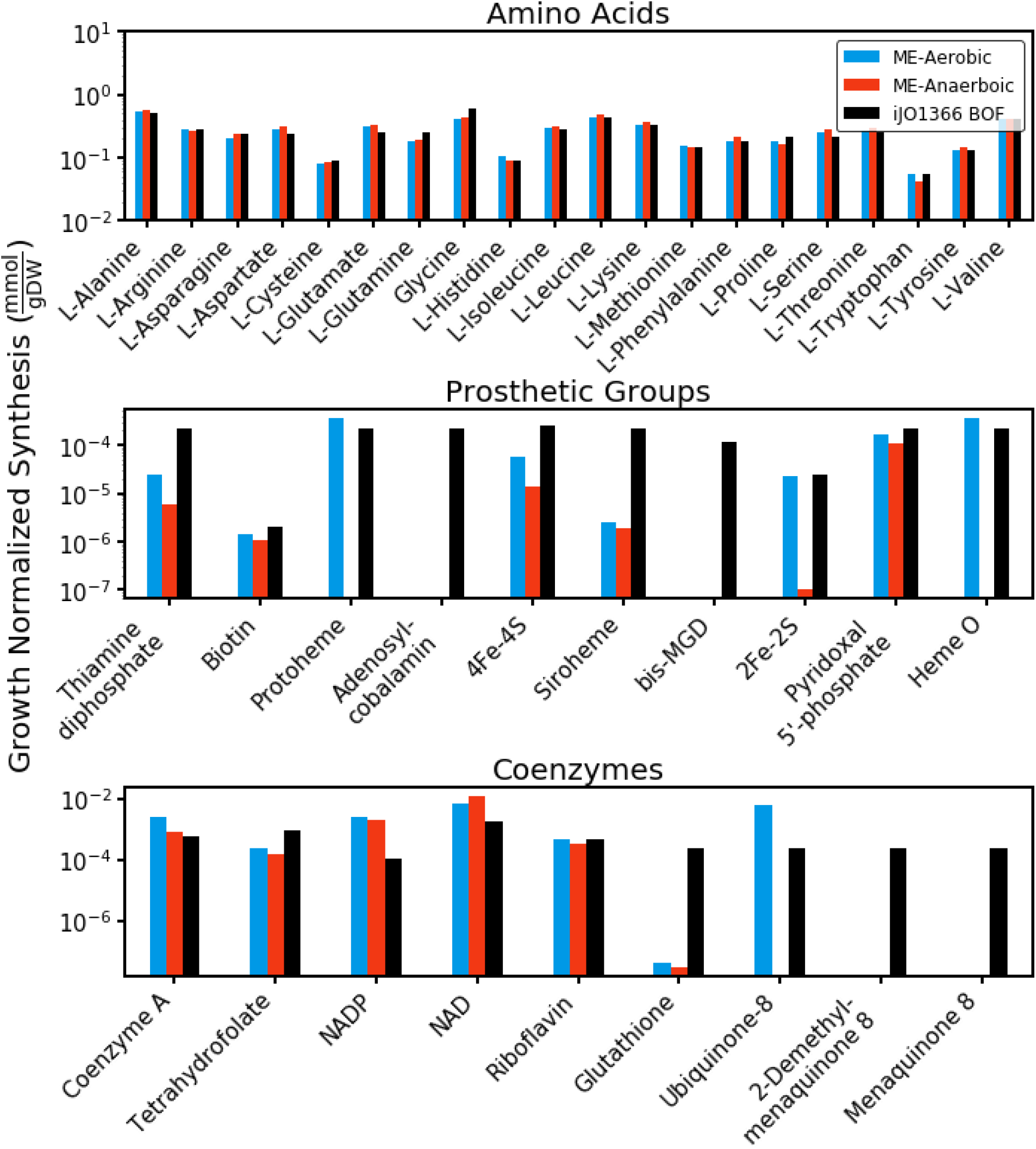
Comparison of ME- and M-model predicted amino acid and cofactor growth-normalized synthesize rates. The ME-model predictions are a function of the predicted intracellular fluxes provided by the simulation, whereas the M-model values are provided by the biomass objective. ME-model predictions are shown for aerobic and anaerobic *in silico* conditions.

Unlike amino acids, many of the cofactor requirements in the iJO1366 BOF are not derived from empirical data. In many cases, they are simply included with small coefficient values to ensure that the activity of the essential gene functions in the cofactor biosynthetic pathways are represented in M-model solutions [5,10]. Therefore, the *quantitative* comparison of the ME-model predicted cofactor usage to the M-model biomass objective function does not provide a high-confidence validation. This comparison does, however, confirm that the ME-model predicts the synthesis of most cofactors within a reasonable range, which is suitable for this study.

The cofactor composition predictions provided by the ME-model are dependent on both the activity of specific reactions in a computed solution as well as the kinetic parameters used to couple reaction flux to enzyme abundance [18]. Thus, a stark difference in cofactor demand is expected when comparing two different computed metabolic states, which is observed for aerobic and anaerobic states (**Figure 2**). Reactions that are less utilized in anaerobic conditions, such as oxidative phosphorylation reactions and pyruvate dehydrogenase, see a decrease in their accompanying cofactors, ubiquinone and thiamine diphosphate (vitamin B1), respectively.

There are a few discrepancies between the *i*JO1366 BOF and ME-model predicted biomass compositions. For example, glutathione is predicted to be synthesized at a rate far lower than the iJO1366 BOF suggests. This discrepancy is unsurprising given that a primary function of glutathione in biological systems is to detoxify the cell in response to external and endogenously produced reactive oxygen species (ROS) stress [19]. Endogenous sources of ROS stress are not within the scope of this model and thus most glutathione activity is not captured. Extensions to *i*JL1678b that explicitly account for ROS damage and detox would more accurately account for the use of glutathione [20].

#### (2) Growth condition-dependent biomass composition

The modified *i*JL1678b model was used to simulate growth on 557 nitrogen, phosphorus, sulfur, and carbon sources under aerobic and anaerobic *in silico* conditions (**S1 Data**). The computed demand of each cofactor and amino acid was normalized by the computed growth rate to allow a direct comparison of the simulations across different growth conditions (see **Methods**). The computed micronutrient demands from the 592 growth-supporting simulations effectively provided condition-dependent biomass objective functions predicted *de novo* from the ME-model. The differences in cofactor and amino acid demand stem from the fact that the simulation for each feasible growth supporting nutrient possesses a unique metabolic state sustained by a unique proteome. Thus, the cofactors and amino acids required to support this proteome differ accordingly.

A high degree of variability was observed for many of the micronutrients depending on nutrient source and aerobicity. This variability was particularly notable for the cofactors which were found to vary in most cases by several orders of magnitude (**Figure 3A**, using BIGG IDs listed shown in **Table B in S1 Appendix**). The range of amino acid demands across conditions, however, was much more narrow. The standard deviation of each micronutrient was further observed as a function of nutrient source and aerobicity (**Figure 3B**). These computations showed that the variation in the cofactor use was largely driven by the carbon and nitrogen sources used for the simulations. Carbon and nitrogen sources also displayed greater variation in the computed growth rates (**Figure A in S1 Appendix**). Little variation in biomass demand and growth rate was observed among phosphorus and sulfur sources. Furthermore, **Figure 3B** shows that the biomass composition variation was similar for nutrient sources in aerobic and anaerobic conditions.

**Figure 3.**
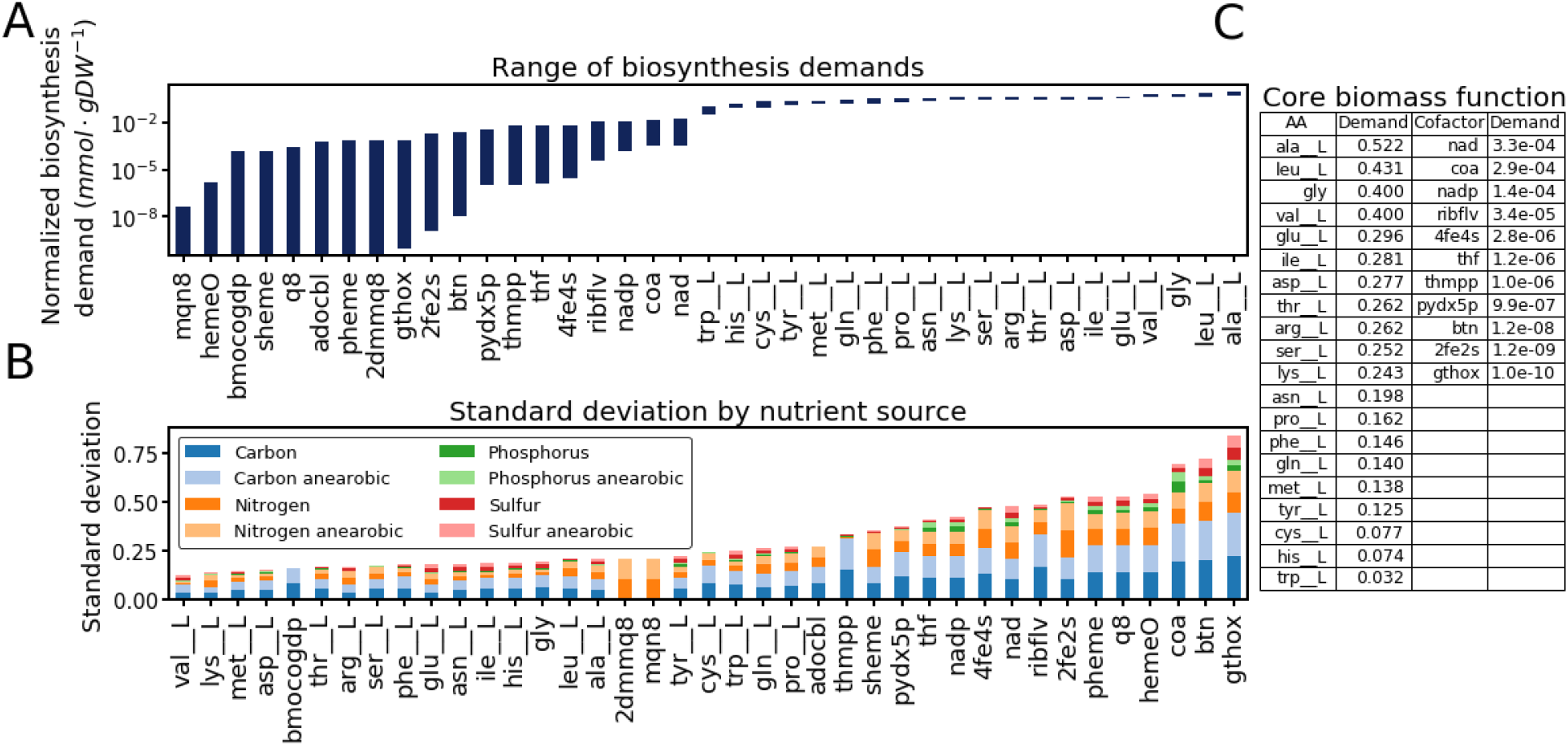
Variation in the synthesis demand of enzyme cofactors and amino acids by growth condition. **A)** The maximum and minimum biosynthesis demand for each amino acid and cofactor. **B)** Stacked bar chart of the standard deviation in the normalized demand for each nutrient source and aerobicity. **C)** The ME-model predicted “core” biomass objective function which shows the minimum demand value for all amino acids and cofactors across the conditions.

The 557 simulated conditions encompass many of the known nutrients capable of sustaining growth in *E. coli* and therefore can be used to infer a “core” biomass composition (**Figure 3C**). The core BOF gives the minimum demand of each biomass component across all of the conditions simulated. This core BOF can be interpreted to imply that, for any given *E. coli* growth condition, the nutrients in **Figure 3C** will need to be synthesized at a rate greater than or equal to the value listed.

Furthermore, a subset of the enzyme cofactors were computationally predicted to be required only in specific growth conditions. The use of some of these cofactors (e.g., ubiquinone-8, protoheme, and heme O) differed based on the aerobicity of the model simulations (**Figure B in S1 Appendix**). This behavior is expected for these cofactors, as they are primarily required for aerobic respiration functions. The demand of other cofactors such as adenosylcobalamin and siroheme are specific to individual growth conditions. For example, adenosyl-cobalamin (vitamin B12) is computationally required for growth only when the carbon or nitrogen source is ethanolamine, since adenosyl-cobalamin is an essential cofactor for ethanolamine ammonia-lyase, the first step of ethanolamine catabolism. Siroheme is computationally required in most growth conditions as a prosthetic group for sulfite reductase, an essential step in the reduction of sulfate to hydrogen sulfide for sulfur assimilation. Growth on other sulfur sources such as cysteine and cysteine derivatives is computationally predicted to alleviate the need for siroheme.

##### Characterizing the aerobic and anaerobic growth by predicted biomass composition

The underlying differences in biomass composition between aerobic and anaerobic growth conditions were also analyzed. PCA decomposition of the computed micronutrient demands showed that aerobic (filled points) and anaerobic (outlined points) simulation could be differentiated along principal components 1 and 2 (**Figure 4**). Further investigation of the vector weightings of principle component 1, showed a general decrease in cofactor use, along with an increase in amino acid use, for anaerobic growth relative to aerobic growth. The most highly weighted amino acids in principal component 1 display an apparent enrichment in prebiotic amino acids [21]. The distinct increase in the use of one prebiotic amino acid, glycine, in anaerobic solutions is shown as a histogram in **Figure 4**. Given that the earliest forms of life existed in a strictly anaerobic environment, it is unsurprising that the enzymes used in anaerobically computed solutions would be enriched in the prebiotic amino acids that were available at the time.

**Figure 4.**
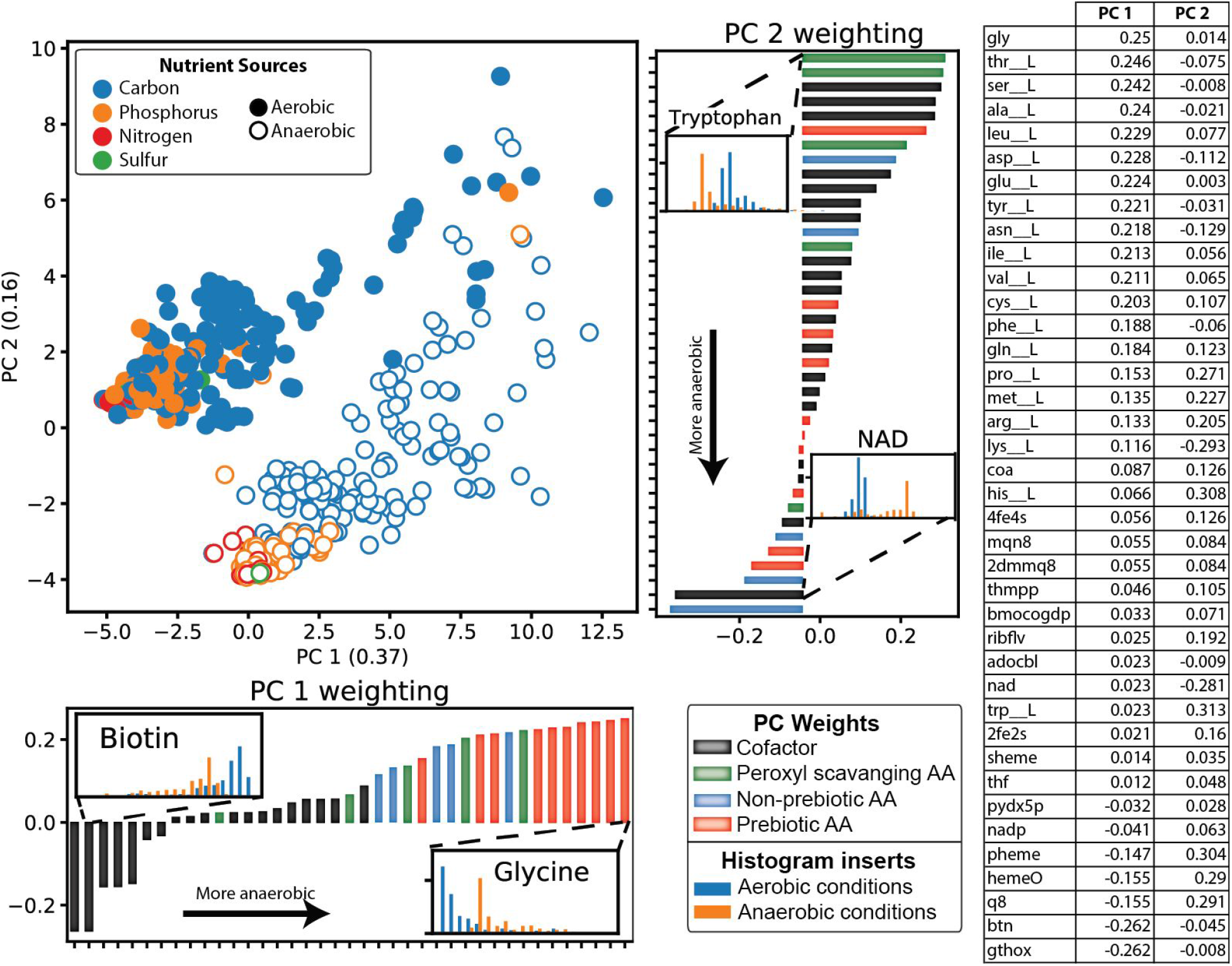
Summary of differences in the synthesis demand of enzyme cofactors and amino acids by aerobicity. PCA analysis of all computed growth conditions reveals aerobic (filled points) and anaerobic (outlined points) growth conditions can be resolved by principal components 1 and 2. Further examination of the component weightings within principal components 1 and 2 can be used to determine the biomass component demands that distinguish aerobic from anaerobic growth. The growth normalized micronutrient demand for two amino acids and two cofactors are shown next to their component weighting. The four histograms demonstrate the clear separation in the aerobicity-dependent demand of these micronutrients. A table of the principal component vector weightings is shown on the right.

Similarly, principal component 2 contributes to the separation between aerobic and anaerobic growth conditions. Though unlike component 1, this component does not show the clear preference of amino acid use in anaerobic conditions and cofactor use in anaerobic conditions. In fact, this component shows that aerobic conditions exhibited increased use of L-histidine and L-tryptophan which are both peroxyl scavenging amino acids, indicating that proteins enriched in these amino acids could be more resistant to oxidative stress. It has been hypothesized that the diversification of amino acids was in part driven by the presence of oxygen and its oxidative properties [22]. Therefore it is unsurprising that aerobic metabolic states would utilize proteins containing these two amino acids.

The component 2 weightings that trended toward anaerobic conditions showed an increase in NAD use. This biomass component was the only cofactor with higher metabolic demand in anaerobic conditions. This observation is expected given the increase in the rate of glycolysis (and thus NAD turnover) observed in fermentative anaerobic metabolism.

##### Clustering growth conditions by predicted biomass composition

Having characterized the major differences in computed biomass compositions during aerobic and anaerobic growth, we next focused on more deeply characterizing differences amongst the aerobic *in silico* growth conditions. It was shown in **Figure 3** that the nutrient source variation was similar for aerobic and anaerobic conditions, and thus only aerobic conditions were used to simplify analysis. The remaining aerobically computed biomass compositions were scaled such that the maximum normalized demand of each cofactor and amino acid had a value of 1 and clustered (see **Methods**). Using hierarchical clustering the 327 growth supporting conditions were effectively partitioned into 6 groups based on the similarity of their micronutrient use (**Table 2**). The samples were displayed on a tSNE plot and colored based on the cluster number (**Figure 5A**). tSNE is a dimensionality reduction technique well-suited for finding subpopulations in high dimensional data. Thus, the higher number of subpopulations seen in the plot suggests that there is some degree of heterogeneity in the biomass composition across the growth conditions. The differences among the nutrient sources could be examined in greater detail if the number of clusters was further increased, but, for ease of interpretation, 6 clusters were used for this analysis (see **Methods**).

**Figure 5.**
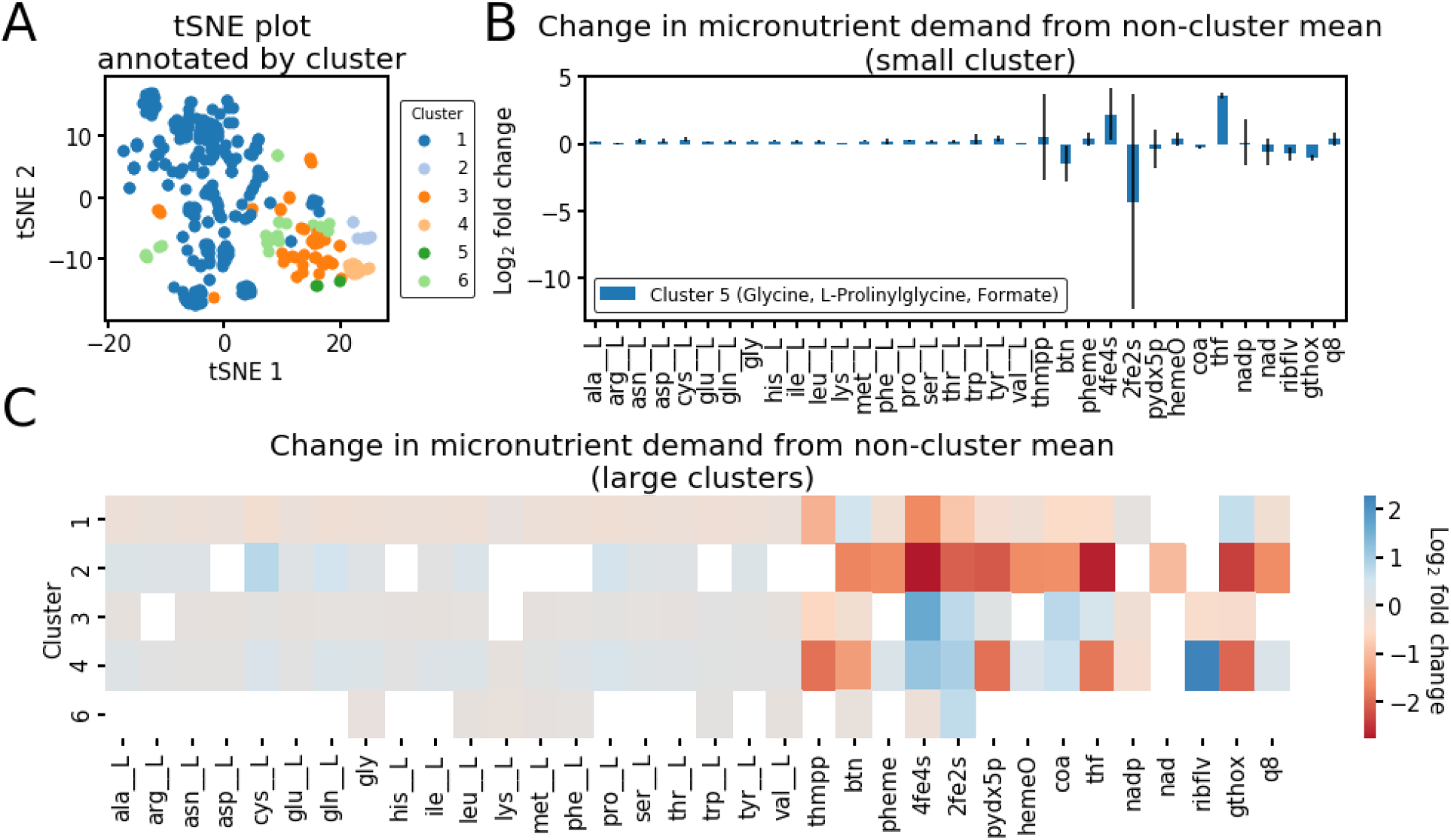
Characterization of aerobic condition-dependent biomass compositions. Hierarchical clustering using Ward’s linkage was performed to divide the *de novo* predicted biomass objective functions into 6 clusters. A) A tSNE plot of the condition-dependent biomass compositions colored by cluster. B) One cluster (cluster 5) contained too few metabolites for meaningful statistical analysis, thus the log_2_ fold change in micronutrient demand compared to the average non-cluster micronutrient demand is shown. A distinguishing feature of growth on carbon sources in this cluster is the high requirement of folates. C) Five clusters contained multiple micronutrient demands that were statistically different (p<1×10^−5^ by Kolmogorov–Smirnov test) compared to the non-cluster demands. The fold change compared to the non-cluster demands for the statistically different micronutrients are shown.

**Table 2:** Summary of clusters determined by hierarchical clustering of aerobic condition-dependent biomass compositions

| Cluster | Cluster summary | Number of conditions in cluster |
| --- | --- | --- |
| 1 | Glucose + nutrients with minimal impact on biomass composition | 231 |
| 2 | Purines and derivatives | 8 |
| 3 | Compounds with pyruvate catabolic intermediates | 47 |
| 4 | Fatty acids carbon sources | 13 |
| 5 | Glycine, prolanyl glycine, and formate as carbon source | 3 |
| 6 | TCA cycle intermediates and derivatives | 25 |

The 6 clusters range in size from 3 to 231 growth conditions and represent groups of growth conditions that exhibit distinct *in silico* biomass compositions. The biomass compositions for the growth conditions within each of these clusters could, therefore, be considered different enough to necessitate a unique biomass objective function (**S2 Data, Figure C in S1 Appendix**). The smallest cluster is cluster 5 and consists of conditions where glycine, prolinylglycine, or formate are used as a carbon source. The distinguishing factor of these 3 carbon sources is their high requirement for folates (~16 fold higher than the average of all non-cluster conditions, **Figure 5B**). Wild type *E. coli* cannot utilize formate or glycine as sole carbon sources, but *E. coli* growth using formate has been engineered by overexpressing formate-tetrahydrofolate ligase along with other components of the folate and serine cycles of *Methylobacterium extorquens AM1*. Further adaptive laboratory evolution of this strain showed that mutations in the rate-limiting step of folate biosynthesis was necessary for robust growth, presumably to increase the rate of folate synthesis [23]. For growth on glycine, the model suggests that this would require a notable increase in the activity of the folate-dependent glycine cleavage system, which could potentially be obtained through adaptive laboratory evolution.

The 5 remaining large clusters were characterized by performing a Kilmogrov-Smirnoff test on the micronutrient demands of the conditions in the cluster compared to all conditions outside of the cluster. The significantly different (p<1×10^−5^) biomass demands for each cluster are summarized as a heatmap in **Figure 5C**. The biomass demands that do not meet this threshold are left blank on the heatmap. This analysis predicted that cluster 2 (purine nutrient sources and their derivatives) are distinct in their reduced requirement of folates, biotin, glutathione, and 4FE-4S iron-sulfur cluster cofactors. Also of note is cluster 4 (fatty acid carbon sources) which is distinguished by the high demand for riboflavin and low demand for folate, pyridoxal phosphate, and thiamine diphosphate needed for growth on these substrates. Wild-type *E. coli* K-12 MG1655 is capable of growing only on long-chain fatty acids, though *E. coli* mutants exist that are capable of growing on short and medium-chain fatty acids as well [24]. This is largely dependent on the strain’s basal expression of β-oxidation enzymes.

The largest cluster was cluster 1 which consisted of all of the growth substrates that are computationally metabolized with proteomes similar to the default growth environment (i.e., glucose M9 *in silico* media). This cluster represented the growth substrates that are predicted to have a minimal impact on the computed biomass composition relative to the default growth state. Cluster 1 included the vast majority of all of the phosphorus and sulfur sources, consistent with the observation in Figure 3B that these sources cause little variability in the biomass composition. These are the conditions in which a biomass objective function obtained from growth on glucose M9 minimal media would be most applicable. However, the subpopulations observed in the tSNE plot in Figure 5A suggests that there is still some degree of variability in the computed biomass compositions within this cluster.

#### (3) Multi-scale analysis of auxotrophy in *E. coli* metabolism

As demonstrated above, ME-models offer the unique ability to comprehensively study how the metabolic needs of an organism can directly influence its use of essential biomass constituents. Alternatively, ME-models can be applied to understand the opposite relationship: the relationship between biomass constituent availability and the metabolic state of the organism. These biomass components, particularly cofactors, are involved in many important cellular functions, thus their activity can have a profound impact on the cellular phenotype [25,26].

Despite the importance of these biomass components for a cell’s metabolic sustainability, many strains of *E. coli* have lost the ability to synthesize some of these cofactors and amino acids throughout their evolutionary history [27]. The evolution of auxotrophy is commonly observed in clinical strains of *E. coli,* and thus understanding the metabolic consequences of auxotrophy can lead to a better understanding of the interactions between pathogenic microbes and their host environment [28,29]. The ME-model was applied to study *E. coli* auxotrophs under *in silico* conditions where availability of the essential metabolite is limited. The predicted cellular response was studied on three levels of resolutions: a phenotypic level, a subsystem level, and by observing changes in the activity of individual reactions.

### Auxotrophy in nutrient limitation

No microbe’s growth environment is constant throughout its lifespan. The same is true for any auxotrophic organism, meaning that auxotrophs will experience periods when the essential nutrients are in excess (**Figure D in S1 Appendix**) and when they are limited. To gauge how nutrient limitation impacts growth, optimal growth rates were computed with *in silico* essential metabolite exposures ranging from the computationally optimal uptake to 1/20th of the optimal value. The *in silico* growth rate was observed with varying uptake rates of an essential amino acid or cofactor. While amino acid limitation elicited a consistent response in the computed growth rate, there was notable variability in the response of the *in silico* cells depending on the essential cofactor (**Figures E and F in S1 Appendix**). Most notably, tetrahydrofolate auxotrophs were computationally predicted to be particularly growth sensitive to drops in tetrahydrofolate availability below the optimal amount (**Figure 6**). This drop in growth occurred in three phases, one in which growth drops gradually to 55.5% of the maximum as folate availability decreased to 65.0% of the optimal. The second phase displayed a sharp decrease in growth from 55.5% to 28.9% of the maximum growth as folate availability decreased from 65.0% to 60.0%. The third phase displayed a gradual decrease in the computed growth rate to 0.

**Figure 6:**
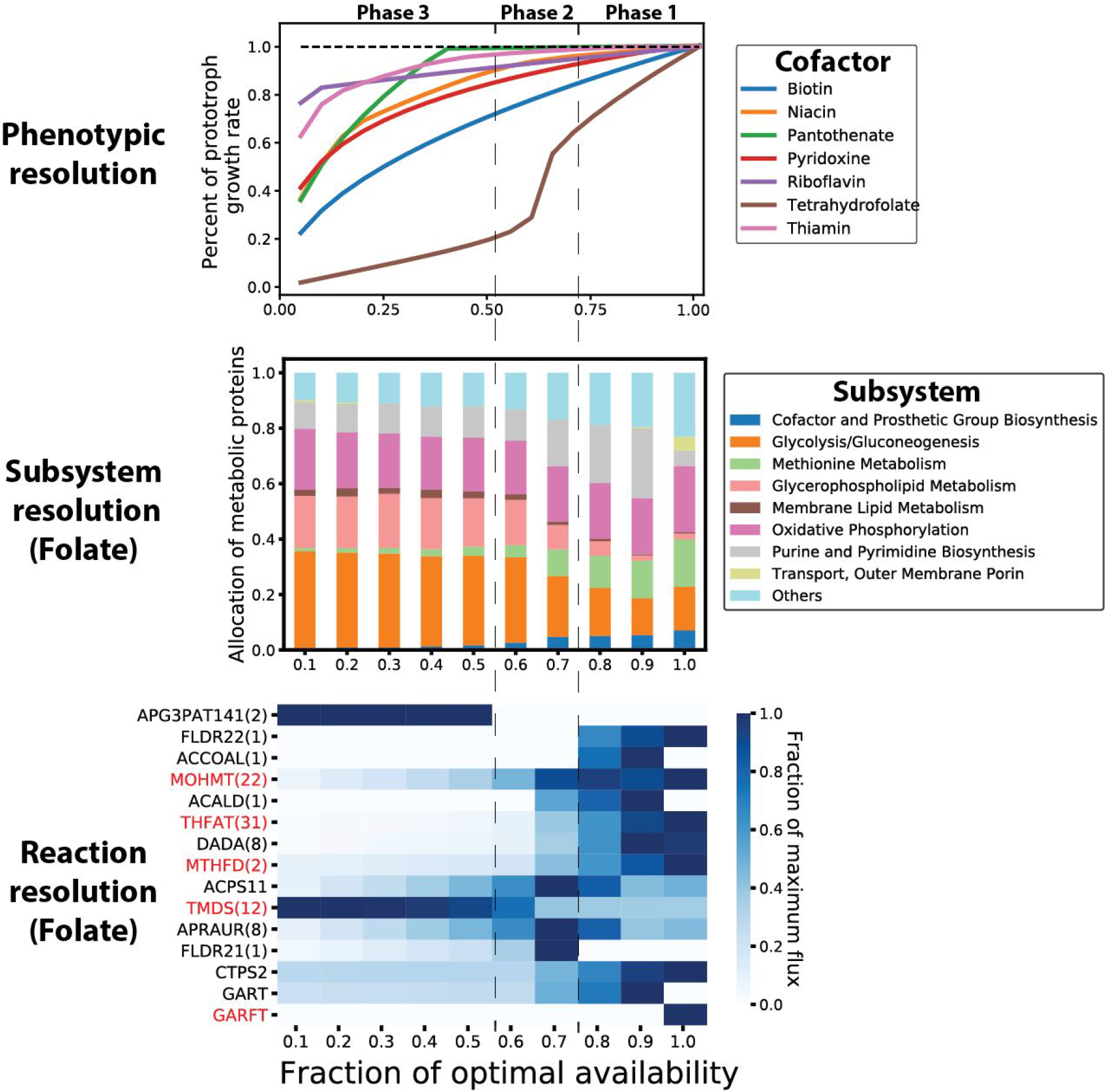
Computed growth rate in auxotrophic models of *i*JL1678b when the availability of the essential cofactor in the legend is limited. For each of the 7 metabolites shown in the legend, reactions were imposed into the model creating a specific auxotrophy for that metabolite. **Top panel**: The growth rate is plotted as a function of the availability of the metabolite indicated in the legend. The percent change in the growth rate compared to the default prototroph model is shown. **Middle panel**: Model-predicted metabolic changes in response to tetrahydrofolate limitation. The fraction of protein allocated to each metabolic subsystem by mass for varying tetrahydrofolate availability is shown (columns) **Bottom panel**: Heatmap showing fraction of maximum growth rate-normalized reaction fluxes. The 15 reactions with the highest standard deviation are shown and are highlighted in red if the reaction relies on folate activity. If reaction fluxes were perfectly correlated throughout the tetrahydrofolate limitation simulations, then these reactions were grouped together. The number in parentheses shows the number of other reactions represented by the row. The right column depicts a simulation with the highest folate availability and the left column depicts a simulation with the lowest folate availability.

The change in protein allocation underlying folate limitation and the resulting drop in growth rate was also assessed. In the first phase of folate limitation, the protein allocated to various subsystems was highly variable, with the amount of protein allocated to purine and pyrimidine metabolism notably increasing. During the drop in growth in phase 2, protein allocation to methionine metabolism, cofactor biosynthesis, and purine and pyrimidine biosynthesis dropped and was reallocated to glycolysis and glycerophospholipid metabolism. The third phase was marked by relatively consistent protein allocation with a minor increase in methionine metabolism and cofactor biosynthesis. The ME-model thus shows a disruption (and thus a compensation by increasing protein allocation) in purine and methionine metabolism, which is known to occur upon treatment with antifolate antibiotics [30,31].

Using the ME-model, metabolic changes can be observed at a higher resolution by characterizing the individual reactions driving the global changes described above (**Figure 6**). For example, when folate is optimally available, the phosphoribosylglycinamide formyltransferase (GARFT) reaction is active which uses formyl-tetrahydrofolate to produce N2-Formyl-N1-(5-phospho-D-ribosyl)glycinamide (fgam), an important intermediate in purine biosynthesis. Folate limitation results in a transition to using GAR transformylase-T (GART), an ATP driven reaction that can produce fgam from free formate. The shift from using GARFT to GART likely contributes to the increase in protein allocation needed for purine metabolism. Additionally, the large drop in growth observed in phase 2 of **Figure 6** coincides with a decrease in 3-methyl-2-oxobutanoate hydroxymethyltransferase (MOHMT), an essential step in Coenzyme A biosynthesis.

Alternatively, a similar analysis was performed for niacin limited growth states (**Figures G and H in S1 Appendix**). In a niacin limited environment, the cellular pools of reduced and oxidized NADP and NAD would be highly depleted. Therefore, an optimally growing cell in this state would likely have to redirect flux into pathways that maximize growth while requiring less of these two cofactors. *i*JL1678b predicts that the optimal approach to optimize NAD and NADP use is: 1) to upregulate the Entner–Doudoroff pathway (bypassing lower glycolysis) 2) to increase activity of the glyoxylate shunt to donate electrons to the quinate pool via malate and 3) to donate electrons to the quinate pool via formate (pyruvate formate lyase (PFL) and formate dehydrogenase (FDH4pp)) and lactate (D-lactate dehydrogenase (LDH_D2)). It is unclear the metabolic route to donate electrons from formate is feasible since PFL is typically only expressed in anaerobic conditions in *E. coli* K-12 MG1655 [32]. Given the rise in antimicrobial resistance to some antifolates antibiotics, understanding metabolic strategies for tolerating nutrient stress could provide clues for how to combat this resistance through possible combinatorial therapies or to potentiate the antibiotic’s activity.

## Conclusions

The presented work provides the first computational study to highlight the systems-level interplay between *E. coli*’s condition-dependent metabolic state and biomass composition. The ME-model of *E. coli* K-12 MG1655 is uniquely suited to examine this relationship, given that it inherently provides predictions of the functional proteome required to sustain a particular metabolic phenotype, including cofactor activity. Using the *i*JL1678b model, simulations were performed on all growth-supporting nutrients in the model, demonstrating notable variability in the demand of cofactors based on the specific growth environment. To determine the metabolic consequences of acquired auxotrophy, the relationship between metabolism and cofactor or amino acid availability was examined. These results provide insight into the metabolic deficiencies that could accompany a drop in the availability of essential nutrients in auxotrophs.

After validating the ME-model predicted biomass objective functions (**Figure 2**), simulations were performed for growth on 557 nutrient conditions aerobically and anaerobically, thus providing 592 *de novo* predictions of condition-dependent biomass objective functions. Analysis of these biomass compositions suggested that unique biomass functions could be appropriate for anaerobic and aerobic simulations. For example, an anaerobic objective function could include an increase in the abundance of prebiotic amino acids such as glycine, along with a decrease in the majority of enzyme cofactors, with NAD being the exception (**Figure 4**). Clustering of the aerobic biomass compositions indicated that the default objective function is suitable for 231 of the aerobic growth conditions (Cluster 1, **Table 2**). Fatty acids, purine and their derivatives, compounds with pyruvate intermediates, and TCA cycle intermediates, however, could potentially benefit from using a modified biomass function. For example, fatty acids could require a biomass objective function deplete in iron-sulfur clusters, biotin, pyridoxal phosphate, heme, folate, NAD, glutathione, and Ubiquinone-8 (**Figure 5**).

ME-models can predict new methods for improving the efficacy of antibiotics either by manipulating the microenvironment of the bacteria or suggesting combinatorial drug therapies. Given that B vitamins provide good targets for antibiotic treatments, the growth condition-dependent cofactor demand could be useful for designing cellular microenvironments to increase or decrease the susceptibility of *E. coli* to some antimicrobials. For example, antifolates that inhibit the synthesis of tetrahydrofolate are a commonly used antibiotic. The analysis in **Figure 5** suggests that treating *E. coli* with antifolates in a glycine-rich environment could potentiate the effect of the antibiotics by increasing the cellular demand of folates. Furthermore, it has been demonstrated that the efficacy of antibiotics can be potentiated by increasing ROS production in *E. coli [33].* Likewise, the predicted metabolic response to cofactor limitation in auxotrophic strains could suggest combinatorial therapies for antimicrobials that target the production of these cofactors. The results in **Figure 6** predict that treating *E. coli* with an antifolate along with an inhibitor of GAR transformylase-T would improve the efficacy of the antibiotic.

This work provides a new look into the inherent coupling between small molecule cofactors and metabolism. A separate integral part of a functional proteome is the metal ion cofactors that form the enzymatic center of many enzymes. Due to the interchangeability of some ion cofactors and intricate mechanisms underlying enzyme mismetallation, fully examining metal ion cofactors was out of the scope of this study [34]. Future work is warranted to exclusively study the metalloproteome and how metal ion availability shapes metabolism.

Lastly, the predictions from this computational study are well suited for future experimental validation. First, many of the cofactor and amino acid auxotrophs are the product of only single gene knockouts in *E. coli* K-12 MG1655, meaning these strains either already exist in single knockout libraries [35] or can be easily synthesized. Adaptive laboratory evolution of these auxotrophs in low concentrations of their essential nutrients could provide valuable insight into mechanisms of *E. coli* adaptation to re-invest its protein toward pathways that maximize growth, while minimizing cofactor use. Second, this work provides predictions of ways to manipulate the growth environment of *E. coli* to potentiate the effect of antibiotics. Future testing these predictions *in vivo* would further underscore the utility of this modeling method.

## Materials and Methods

### Software

All constraint-based modeling analyses were performed using Python 3.6 and the COBRApy software [36]. ME-model operations were performed using the COBRAme framework [16]. Due to the fact that ME-models are ill-scaled [12], qMINOS [37,38], which supports quad (128-bit) precision, was used for each ME-models simulations. M-model simulations were performed using the iJO1366 model of *E. coli* K-12 MG1655 metabolism [17], since *i*JL1678b-ME was reconstructed using this M-model of *E. coli* as a scaffold. All M-model optimizations were performed using the Gurobi (Gurobi Optimization, Inc., Houston, TX) linear programming (LP) solver.

### ME-model modifications for modeling cofactor activity

The *i*JL1678b ME-model of *E. coli* K-12 MG1655 was used for all simulations in this study. The activity of enzyme prosthetic groups are inherent in the ME-model formulation [12], which uses coupling constraints to connect the synthesis of individual enzymes (including their accessory groups) to the reactions they catalyze. Coenzymes (NAD, folates, etc.) have some of the same properties of enzymes in that they are recycled within the network in both M- and ME-models. These models therefore ensure that the coenzymes are balanced, but they do not account for the biosynthesis of these coenzymes. As a result, models have incorporated these coenzymes in the biomass objective function to force their biosynthesis in a way that is independent of their use in the model.

The *i*JL1678b ME-model was thus modified to couple the biosynthesis of coenzymes to their activity, similar to other enzymes in the model. This is accomplished using a pseudo-kinetic term to relate the concentration of the coenzyme pool to its activity throughout the metabolic network, which we will simply call k_activity_. This term represents a very rough estimation of the first-order kinetics of the reactions involving the coenzyme in the network. This term was chosen as 1×10^4^ hr ^−1^ and applied to each reaction where the uncharged version of the coenzyme acts as a reactant:

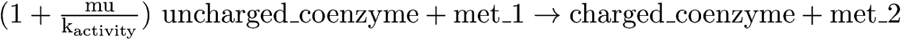

For this study, we were interested in the relative activity of these coenzymes across varying growth conditions. It was thus important that the computed coenzyme abundances were within a reasonable range (**Figure 2**), but the quantitative accuracy of the abundance predictions was not necessary. Therefore, accounting for the complex kinetics of the coenzymes throughout the reconstruction was outside the scope of this work. This simple approach effectively relates the rate of coenzyme biosynthesis with its metabolic activity and growth rate.

Changes and corrections were applied to *i*JL1678b-ME to allow the model to be used for this study. First, the biomass constituent demand reaction flux was set to zero. This reaction is included in the default ME-model to account for the synthesis of many of the coenzymes whose activity is modeled directly with the modified ME-model. It is thus no longer necessary. Furthermore, the uptake of any metal cation not included in M9 media was constrained to zero. Malate oxidase was made irreversible as suggested in Monk *et al.* [39]. Corrections were further made to *i*JL1678b-ME to more accurately compute prosthetic group use. For example, acetolactate synthase (ilvH and ilvI) and 2-oxoglutarate dehydrogenase (sucA, sucB, and lpd) were updated to correctly require FAD and thiamine diphosphate as prosthetic groups.

### ME-model parameterization and optimization procedure

The k_eff_ coupling parameters [12] for each metabolic reaction in *i*JL1678b-ME were determined based on a machine learning approach that incorporated enzyme features, network properties, and proteomics data to predict k_eff_s [18,40] from a set of *in vivo* derived enzyme turnover rates [41]. In order to capture the high catalytic efficiency and encourage model activity of pyruvate dehydrogenase the k_eff_ of this reaction was set to 1500 s^−1^ [42]. The remaining k_eff_s for expression machinery and transport reactions were set to a default value of 65 s^−1^ [12], as these processes were out of the scope of the machine learning approach. The unmodeled protein value [12] was set to 0 for all simulations. All remaining parameters were set to their default values [16].

Due to non-linearities stemming from the enzyme coupling constraints, ME-models cannot be optimized directly as an LP and thus are solved using a binary search algorithm. To perform the binary search, the following procedure was implemented. First, each symbolic coefficient (growth rate, μ) or reaction bound was compiled into a function by sympy [43]. Then, an LP file was created for the qMINOS solver with all of these symbolic functions evaluated to 0. While the model will always be feasible at 0, starting with a known feasible point results in a basis that can be used to speed up the next run. Afterward, for each instance of the binary search in μ, values in the LP were replaced by recomputed ones, and the problem was resolved using the last feasible basis. This approach was continued until the maximum feasible μ and minimum infeasible μ were within the defined tolerance (1×10^−13^).

### Computing biomass constituent demand

The amino acid biosynthetic demand was determined based on the translation flux and amino acid composition of each protein in the model. Prosthetic group demand was determined based on the sum of the complex formation fluxes of all enzyme complexes containing the appropriate prosthetic group. Due to the high connectivity of coenzymes throughout the metabolic network, the biosynthetic flux of each cofactor was determined based on the activity of a reaction in its biosynthetic pathway (**Table A in S1 Appendix**). This was sufficient given that each cofactor contains one direct biosynthetic pathway. To enable a direct comparison of the ME-model to the iJO1366 biomass objective function, the computed biomass constituent demand values were normalized by the computed growth rate.

### Clustering analysis

The biosynthetic demand for each micronutrient was normalized by its maximum value across all aerobic simulations. Hierarchical clustering was then performed using ward linkage criteria, as implemented in the scikit-learn Python package. The NbClust and factoextra R packages were used to optimize the number of clusters using the elbow method and silhouette method (**Figure I in S1 Appendix**). The average silhouette width was at a global maximum at 2 clusters, but there were additional local maxima at 6 and 17 clusters. The intra-cluster variation was also computed for each cluster with the intent of finding an elbow in the curve (i.e., a point in which adding additional clusters provides diminishing returns). Since the elbow appears to also occur at a local maxima for silhouette width, a cluster number of 6 was chosen.

### Reproducibility

All of the code necessary to reproduce the presented results can be found on GitHub at https://github.com/coltonlloyd/me_biomass.

## Acknowledgments

We would like to thank Zachary King, Justin Tan, Bin Du, and Joshua Lerman for informative discussions. Funding for this work was provided by the Novo Nordisk Foundation through the Center for Biosustainability at the Technical University of Denmark under Grant NNF10CC1016517 and the NIH National Institute of General Medical Sciences under Grant numbers NIH R01 GM057089 and NIH 1-U01-AI124316. This research used resources of the National Energy Research Scientific Computing Center, which is supported by the Office of Science of the US Department of Energy under Contract No. DE-AC02-05CH11231.

## Supporting information

**S1 Appendix. Supplemental tables and figures.**

**S1 Data. Computed growth rates for all growth conditions.**

**S2 Data. Hierarchical clustering summary.** The growth conditions represented in each cluster along with the average biomass composition.

